# A mathematical model for axonal transport of large cargo vesicles

**DOI:** 10.1101/2023.01.18.524542

**Authors:** Nizhum Rahman, Dietmar B. Oelz

## Abstract

In this study, we consider axonal transport of large cargo vesicles characterised by transient expansion of the axon shaft. Our goal is to formulate a mathematical model which captures the dynamic mechanical interaction of such cargo vesicles with the membrane associated periodic cytoskeletal structure (MPS). It consists of regularly spaced actin rings that are transversal to the longitudinal direction of the axon and involved in the radial contraction of the axon. A system of force balance equations is formulated by which we describe the transversal rings as visco-elastic Kelvin-Voigt elements. In a homogenisation limit, we reformulate the model as a free boundary problem for the interaction of the submembranous MPS with the large vesicle. We derive a non-linear force-velocity relation as a quasi-steady state solution. Computationally we analyse the vesicle size dependence of the transport speed and use an asymptotic approximation to formulate it as a power law that can be tested experimentally.

## 1 Introduction

Neurons are the foundational units of the nervous system. They are involved with information processing and transport along their long cellular extensions termed dendrites and axons. Tightly regulated axonal transport of multiple types of cargo such as synaptic vesicles, organelles and proteins packed into cargo vesicles is paramount for the functioning of the neuron. This involves anterograde axonal transport driven by molecular motors of the kinesin family and dynein-dependent retrograde transport [1, 2].

The axon shaft is characterised by a particular cytoskeletal organisation termed membrane associated periodic skeleton (MPS). It involves a sub-membraneous actin cortex and subcortical evenly spaced actin rings with a periodicity of 180−190nm [3, 4]. They are made of a few long intertwined actin filaments forming actin braids [5]. The MPS stabilises the axon diameter [6–8]. Indeed it has been reported that F-Actin disassociation leads to the collapse of axons [9] whereas inhibition of the F-Actin capping protein *α*-adducin leads to axon diameter expansion [10]. Non-muscle myosin IIa and IIB are associated with the periodic actin rings [6] and involved in radial contractility [11].

The protein spectrin couples neighbouring rings through an elastic connection and contributes to the stability of the axon [6, 12]. A longitudinally aligned system of cytoskeletal elements involves F-actin trails, neurofilaments and microtubules [8]. Particularly the latter serve as tracks for the molecular motors involved in axonal transport.

The diameter of the axon shafts varies among different tissues and species and ranges from 0.1 μm to several microns [13]. Cargo sizes vary within a wide range which includes mitochondria (0.75–3μm) [14], autophagosomes (0.5–1.5μm) [15] and endosomes (50nm–1μm) [16]. Thus, cargo sizes are comparable to the size of various CNS (central nervous system) axons, often exceeding it. Radial axonal contractility in combination with transient expansion of axon calibre facilitates the passage of large cargoes. When a large cargo passes through the axon, the axon acts like an elastic material avoiding the further delay of cargo passage. Subsequent radial contraction of the transversal actin rings is myosin dependent[11]. Yet, whether the actin rings are open loops, or closed rings (see figure 1) is an open question [6].

**Figure 1:**
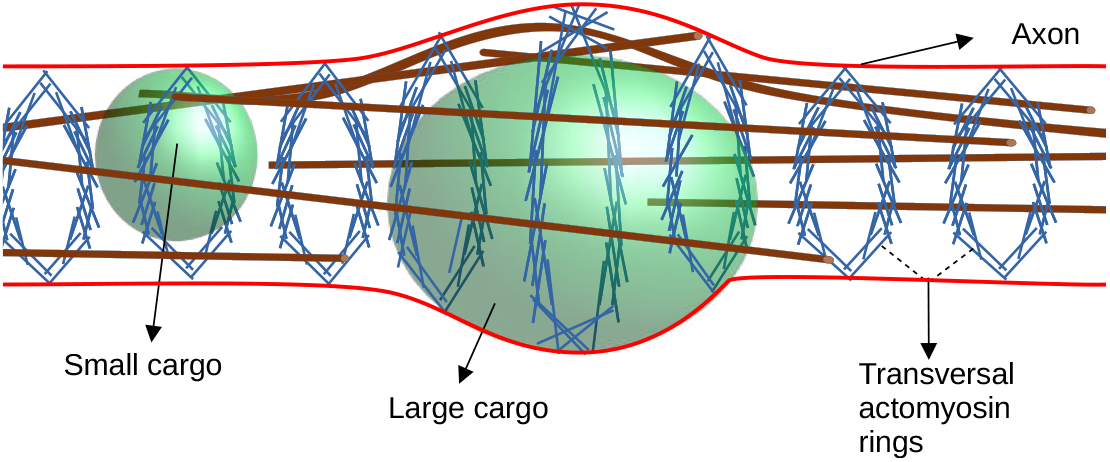
Actin rings (blue, sketched according to the closed ring hypothesis [6]) are wrapped around the circumference of the axon with equidistant spacing. Vesicles of various sizes (green) are moving through the axon. Microtubules (brown) provide transport rails for molecular motor driven transport of vesicles.

The speed of axonal transport varies significantly between slow axonal transport (*<* 0.1 μm*/s*) and fast axonal transport (*>* 0.5 μm*/s*) [17]. For large cargo particles, whose diameter exceeds the diameter of the axon shaft, it has recently been reported that the speed of retrograde axonal transport is inversely correlated with the diameter of the cargo particles, i.e. large-diameter cargo moving slower than smaller-diameter cargo [11].

One possible explanation is size-dependent friction on the cargo originating from the mechanical interaction with the axonal cortex [11]. This hypothesis is also supported by a simlation study suggesting that the effective viscosity between organelles and the axon wall at a small distance can become large enough to impede cargo transport[18].

Another potential mechanism, which might explain the slower speed of transport of large cargo particles, is tight spatial alignment of microtubules in small calibre axons which could stabilise the pool of molecular motors engaged in active transport. When the axon is stretched by large calibre cargo distances between parallel microtubules might be larger and impede likely reattachment of molecular motor proteins after detachment [18].

Despite these theories, it is an open question how axonal radial contractility regulates cargo transport [11]. In particular, the observation that in myosin inhibited cells trajectories of large cargo retrograde transport visualised through kymographs appear more random and even exhibit frequent backtracking [11] seems to contradict the friction hypothesis, but could be supported by the microtubule alignment hypothesis. Another potential explanation could originate from the direct mechanical interaction of the contractile actin rings. In this study, it is our goal to develop the modelling framework to investigate this interaction between the transversal actin rings and large cargo particles.

To this end, we formulate a detailed mathematical model which couples the active and passive mechanical properties of the actin rings modelled as visco-elastic Kelvin-Voigt elements. We start by formulating the model in a time-discrete setting with a fixed spacing of the actin rings. In a second step, we formally pass to a continuum limit of both time and actin ring spacing, both in section 2. In the following sections, we process the model further. In section 3 we non-dimensionalise. The resulting mathematical model resembles the classical obstacle problem [19] which is a model problem in the study of free boundary problems. In this study, the free boundary is the interface where the cargo vesicle touches the axonal cortex. We consider quasi-stationary solutions of this problem from which we derive the force-velocity relation for the transport of large cargos in section 4. In section 5 we study the dependence of transport speed on vesicle size numerically, and we use asymptotic approximation to formulate the decrease of transport speed for large vesicle sizes as a power law valid for vesicle sizes which are close to the axon size.

## 2 Mathematical Model

Our goal is to develop a mathematical model which relates the forces, especially those exerted by the transversal actin rings and their associated myosin motor proteins, to the velocity of large cargo vesicles undergoing axonal transport.

To introduce the model we idealise the axon as a one-dimensional structure decorated with concentric rings located at equi-distant positions *y*_*j*_ ∈ ℝ (figures 1 and 2). Here we use *j* = 1…*M* as an index for actin rings assuming – for notational convenience – the presence of *M* actin rings in total.

**Figure 2:**
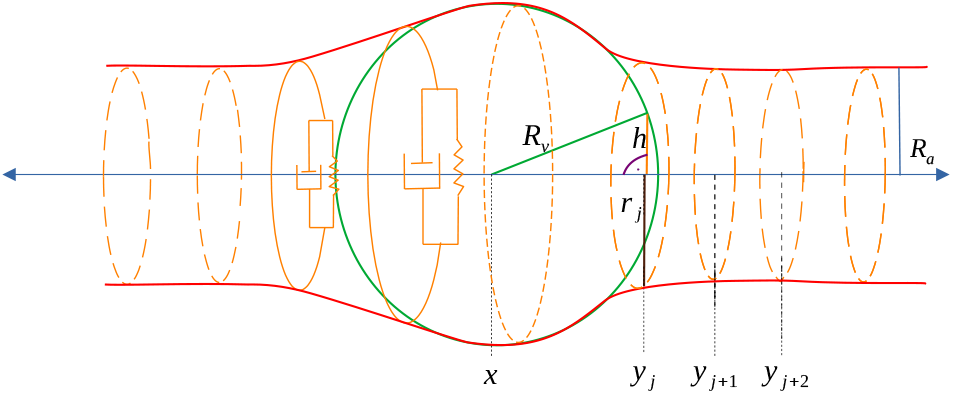
Sketch of the constituents of the mathematical model depicting a cargo vesicle at position *x* ∈ ℝ (green with radius *R*_*v*_) moving from left to right through the axon with radius *R*_*a*_. As it is moving through the axon it extends the radii *r*_*j*_ of the actin rings uniformly positioned at *y*_*j*_, *y*_*j*+1_, etc. The function *h* at every point denotes the “height” of the vesicle.

The time-dependent degrees of freedom of the model are the position *x* = *x*(*t*) ∈ ℝ of a large cargo vesicle with radius *R*_*v*_ and the radii of actin rings written as *r*_*j*_ = *r*_*j*_(*t*) *>* 0 where *t* ≥ 0 refers to time.

As a result of our lack of more detailed knowledge about the mechanical properties of transversal actomyosin rings and in order to keep the model amenable to mathematical manipulations in the final sections of this study, we will usually assume linear expressions for forces, i.e. square energies, and we also assume that the cargo vesicle is a solid spherical object rather than a viscoelastic, structurally inhomogeneous object. This should be rather seen as a modelling assumption and not as an assumption of inifinitesimal strains since the strain of actomyosin rings can reach 0.5 [11]. Finally, we neglect the presence of microtutubles (see figure 1) and larger organelles in the axon and idealise the axon-cargo geometry in the sense of figure 2 (figure 2 in [11] shows an electron micrograph of large vesicles and surrounding microtubules).

We construct the model as a system of force-balance equations taking into account the following forces acting on the cargo vesicle and the actin rings.

We assume that the cargo vesicle is pulled by the given force *F*, which, depending on direction, may be positive or negative, and that through its interaction with the cytoplasm the cargo vesicle is exposed to the drag friction *F*_friction_, which we model as linear drag with coefficient *ξ >* 0,

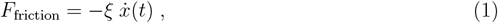

where 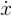 is the notation we use for the velocity of the cargo vesicle. Note that when modelling the pulling force exerted by molecular motors [20, 21] through an affine force-velocity relation, the parameters *F* and *ξ* will be determined by stall force and free moving velocity of the molecular motors.

We model the actin rings as visco-elastic Kelvin-Voigt elements with equilibrium radius *R*_*a*_ (figure 2). This implies that the elastic and viscous forces exerted by every ring are in parallel. Its elastic component, which in (figure 2) is visualised by a spring, is given by

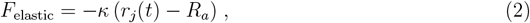

where *κ >* 0 is the constant of elasticity. Note that this implies that we are modelling myosin motors indirectly as drivers of radial contraction, namely through the equilibrium radius of actin rings *R*_*a*_, and through their stretching stiffness *κ*.

The viscous resistance to changes in actin ring size – in figure 2 visualised by a dashpot – is given by

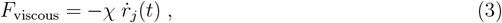

where *χ* is the coefficient of viscosity. It models the internal resistance of the actin rings to shortening and stretching causing stretching and realignment of actin fibers and cross-linker proteins as well as transient unbinding from crosslinker proteins and myosin motors.

We assume that the cargo vesicle and the actin rings are mechanically interacting: when the size of the vesicle is close to the radii of the rings, the vesicle would push the actin rings apart so it can pass. The resulting force is modelled as a reactive force resulting from the impenetrability of vesicles and rings.

Our goal is to formulate the equations of motion as a system of force-balance equations for the conservative and dissipative forces listed above in (1), (2), (3) as well as the external pulling force *F* and the repulsive forces where axon and cargo touch.

To this end we will start in a time-discrete setting, formulate the model first as a minimisation problem for a potential energy functional for a fixed point in time, and then derive the system of force-balance equations as Euler-Lagrange system.

The advantage of this approach which we call *“generalised-gradient-flow”* is that at first we only need to formulate the *scalar-valued* energy functional[22–24]. In the second step, we recover the system of *vector-valued* force-balance equations for both axon and cargo through an analogous application of the least action principle. In the variational equations, once we formally let the size of time steps and scaling of penalising potentials such as (6) converge to zero we recover the force-balance equations consisting of conservative and dissipative forces and Lagrange multipliers in continuous time.

Note that this approach can be interpreted as a formulation of constrained Lagrangian mechanics in the over-damped regime which enables us to neglect kinetic energy. The initial formulation in the time-discrete setting enables us to recover viscous forces from dissipative energies.

Here, to formulate a closed system of equations we introduce the following variational formulation of the model, in which every force term above is represented by a corresponding conservative or dissipative energy, and in which the constraint is represented by a penalising potential. We compute the time-discrete solution 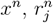 for every index of time *n* = 1, … with timesteps Δ*t* as a minimiser,

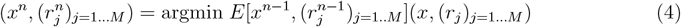

of an energy functional with respect to the configuration of the system, i.e. actin ring radii and vesicle position. The functional is also parametrised by the configuration of the system at the previous point in time written as *x*^*n*−1^, 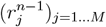. Its components are

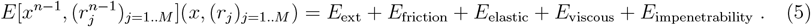

The components of the energy functional corresponding to conservative and dissipative forces are

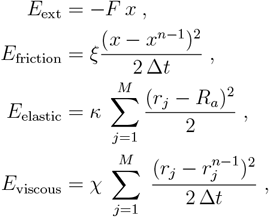

which are chosen in a way such that the negative variation of these energies evaluated at *x*^*n*^, 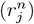) corresponds to the forces listed above. For dissipative forces this holds in the limit when Δ*t* → 0, e.g. 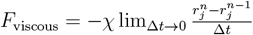.

The penalising potential which enforces the constraint that the actin rings do not penetrate the vesicle is given by

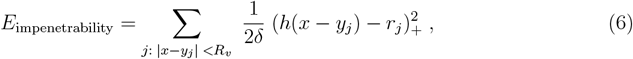

where *δ* is a small parameter implying that the coefficient in front of the penalising potential is large. In (6) we restrict the sum to those actin rings which are sufficiently close for interaction with the vesicle and we use the subscript (.)_+_ as a short notation for the positive part of what is inside the bracket, i.e. (*x*)_+_ = *x* if *x >* 0, otherwise it is zero. The respective term contributes to the sum (6) if the cross-sectional radius *h* of the vesicle at the position of the ring is larger than the radius of that actin ring. It is computed as a function of the distance between ring and vesicle position, *h* = *h*(*x* − *y*_*j*_) according to (see figure 2)

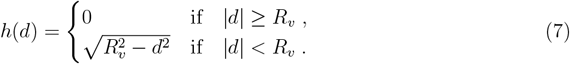

This finalises the formulation of the mathematical model as a variational scheme in the time-discrete setting.

Next we will derive the equations of motion. The solution to the variational scheme is characterised by 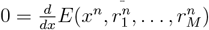, which implies

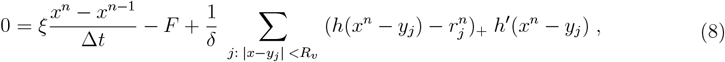

and by 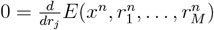 for every *j* = 1 … *M*, which yields

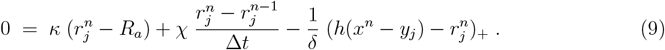

In the formal limit when the time steps are small, i.e. *dt* → 0, we write 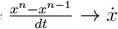 and 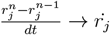 and the system (8), (9) reads

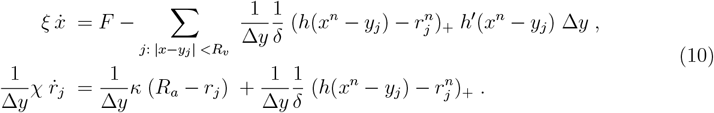

In the formal limit when the penalisation of the impenetrability constraint becomes large and the spacing of the actin rings small, *δ* → 0 and Δ*y* → 0, we define the Lagrange multiplier formally as a limit,

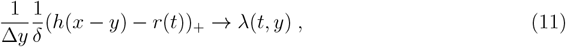

which is a force density corresponding to the interaction force between vesicle and actin rings. We also identify 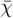 with the limit of *χ/*Δ*y* and 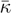 with the limit of *κ/*Δ*y*. Note that the vesicle position *x* = *x*(*t*) is a dependent variable whereas *y* ∈ ℝ is an independent variable of our model. We get the following system of two equations for the functions *x* = *x*(*t*) (vesicle position), *r* = *r*(*t, y*) (actin ring/axon radius) and *λ* = *λ*(*t, y*) (interaction force density), see figure 3,

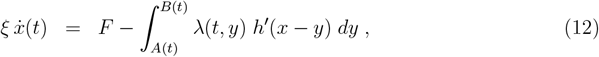

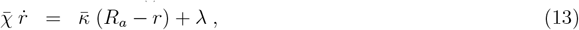

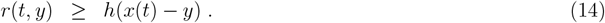

Here *A* and *B* define the interval (*A, B*) where the axon - represented by the transversal actin rings – touches the vesicle, i.e. where (14) is an equality. This equality determines the interaction force density *λ* = *λ*(*t, x*) within the interval (*A, B*) ⊂ ℝ, outside of which *λ* is zero.

**Figure 3:**
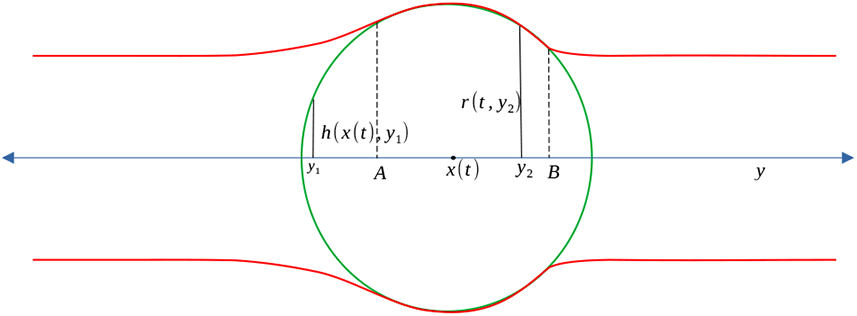
The degrees of freedom of the continuum model (12)-(14): The cargo vesicle at position *x*(*t*) moving from left to right is touching the axon with radius *r* = *r*(*t, y*) along the interval (*A, B*). At every position *y* the function *h*(*x*(*t*), *y*) denotes the “height” of the vesicle, i.e. the radius of the segment of the vesicle which is perpendicular to the axon.

Note that an additional equation is derived as follows: In figure 4 which is sketched in a regime where both time and the spacing of actin rings are discrete, we consider the boundary point at *B* and assume that it moves to the left, i.e. 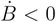. We consider the actin ring at *y*_*j*_ which at time *n* is immediately to the right of *B*^*n*^ and which at the previous point in time *n* − 1 is immediately to the left of the interface point. This suggests that the force balance equation according to (9) is given by

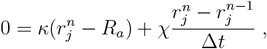

where the penalisation term for the interaction of vesicle and axon is zero. Since at the previous point in time *n* − 1 this actin ring does touch the vesicle, we replace 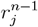 by *h*(*x*^*n*−1^ − *y*_*j*_). In the homogenisation procedure by which Δ*t* → 0 and Δ*y* → 0 we may place the ring *j* arbitrarily close to *B*. By continuity we replace 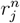 by *r*^*n*^(*B*) = *h*(*x*^*n*^ − *B*) obtaining 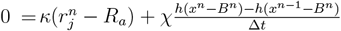. In the aforementioned limit we find that

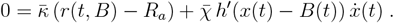

On the other hand, the limit of (13) evaluated at a sequence of points which converges to *B*(*t*) from the right implies that 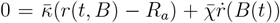. Therefore 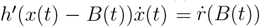.

**Figure 4:**
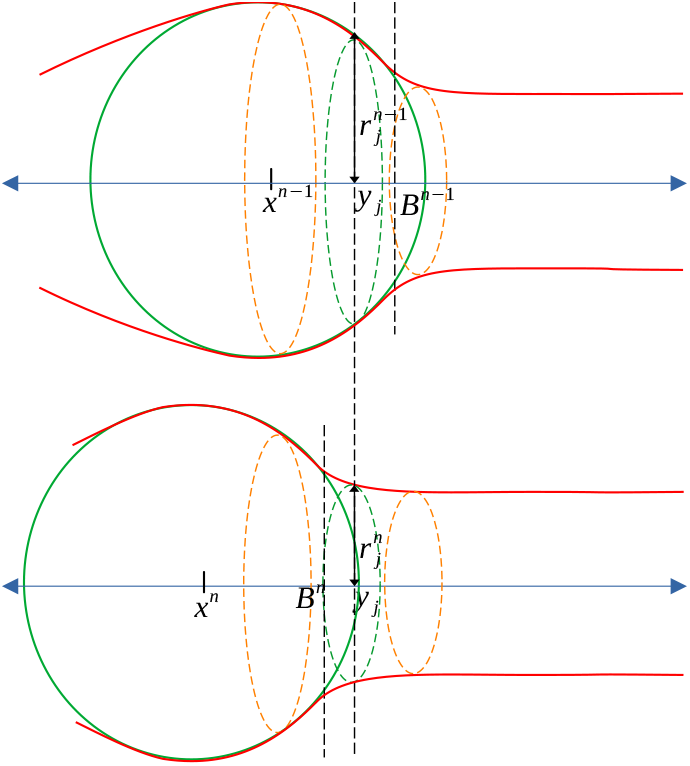
Derivation of the free boundary condition at the rear of the vesicle (here moving from right to left).Comparison of vesicle position and ring sizes at time *n* − 1 (above) and time *n* (below).

Note that the opposite assumption, i.e. 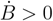, does not imply such a condition. For a fibre that would be immediately left from *B*(*t*), though at any previous point in time right of *B*(*t*), the force balance equation (9) would have a non-zero interaction term. The same equation taken from away of the vesicle would not involve this term 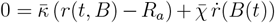 and would therefore not allow to draw a conclusion about the time derivatives.

Finally, the same arguments applied the left interface point assuming that 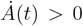 suggests that

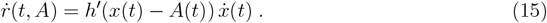

## 3 Nondimensionalization

To facilitate the further analysis of the model (12)-(14) we non-dimensionalise introducing the re-scaled independent and dependent variables 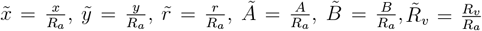 and 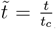, where *t*_*c*_ is the reference scale for time which we will determine further below. The system (12)-(14) becomes

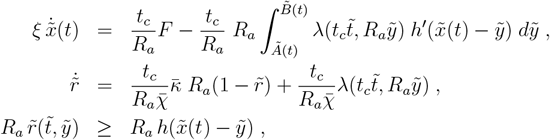

where we used that 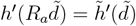 (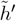 is defined by (7) for the scaled vesicle radius 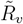). We determine *t*_*c*_ such that 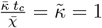, i.e. 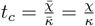. Let 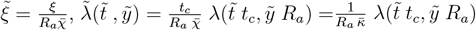 and 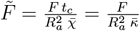. We obtain

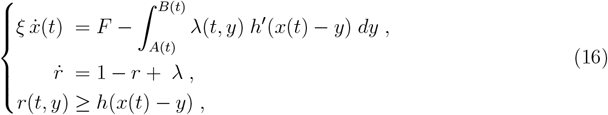

where we have omitted the tildes again. Note that the interaction force vanishes, *λ* ≡ 0, where the axon does not touch the vesicle, i.e. where *r*(*t, y*) *> h*(*x*(*t*) − *y*).

The parameters of the model are listed in table 1. We estimate that the cargo velocity *V* is of the order of magnitude 1.5 μm/sec [11] and therefore we determine the drag friction coefficient as *ξ* ≈ *F/V* = 10 pN sec/μm having in mind that this refers to a cargo vesicle the size of which does not exceed the axon diameter. Furthermore, with the parameters listed in table 1, we obtain the dimensionless parameters 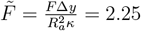 and 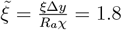.

**Table 1:** List of parameters values

| Description | Symbol | Value | References |
| --- | --- | --- | --- |
| Radius of cargo vesicle | $R_v$ | 0.3 $\mu\text{m}$ | [11] |
| Equilibrium radius of axon | $R_a$ | 0.2 $\mu\text{m}$ | [11, 13] |
| Pulling force | $F$ | 15 pN | estimated |
| Spacing between the actin rings | $\Delta y$ | 0.18 $\mu\text{m}$ | [3] |
| Drag friction | $\xi$ | 10 pN $\mu\text{m}^{-1}$ sec | estimated |
| Stretching stiffness of actin rings | $\kappa$ | 30 pN $\mu\text{m}^{-1}$ | estimated |
| Viscosity of actin rings | $\chi$ | 5 pN $\mu\text{m}^{-1}$ sec | estimated |

## 4 Force-velocity relation

In this section, it is our goal to determine the force-velocity relation predicted by our model as the quasi-stationary solution characterised by a constant vesicle velocity *V* which we assume to satisfy *V >* 0. This implies that *x*(*t*) = *V t* as well as 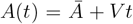 and 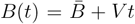, where we expect 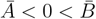 (see figure 5).

**Figure 5:**
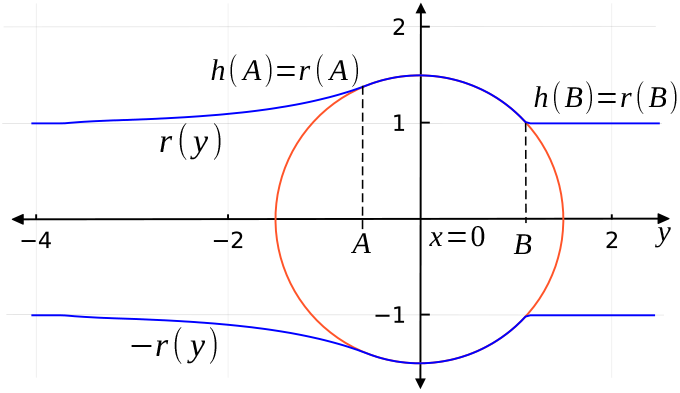
Travelling wave solution with vesicle positioned at *x* = 0 for the dimensionless parameters *F* = 2.25 and *ξ* = 1.8 corresponding to the physical parameters listed in table 1. The cargo (red) moves from left to right and touches the axon (blue) along the interval [*A, B*].

The ring/axon radius *r* as well as the interaction force/Lagrange multiplier *λ* in this scenario will be simply advected with speed *V* and have the structure of travelling wave solutions 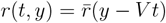 and 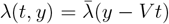, where 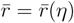 and 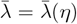 only depend on the travelling wave variable *η* = *y* − *Vt*.

With this ansatz – omitting the bar-notation again – the system (16) becomes

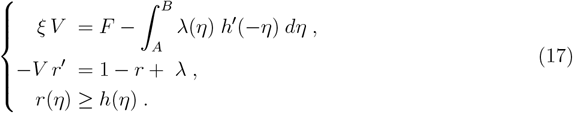

where we used *h*(−*η*) = *h*(*η*). In addition, since 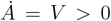, the free boundary condition (15) translated into travelling wave variables holds, i.e. *r*^*’*^(*A*) = *h*^*′*^(*A*). A numerical solution corresponding to the travelling wave solutions is sketched in figure 5.

We solve the equation −*V r*^*′*^ = 1 − *r* on the interval (−∞, *A*) where the axon and the vesicle do not touch and find the general solution *r*(*y*) = *C*_1_ *e*^*y/V*^ + 1.

Now we consider the interface point at *A* where axon and vesicle touch. It therefore holds that *r*(*A*) = *h*(*A*) which allows to infer the value of *C*_1_. We find that

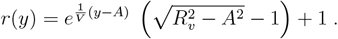

The solution of *r*(*y*) between points *A* and *B* is the height of the vesicle *h*. We make the modelling assumption that as the vesicle moves it progresses into an axon which is initially not stretched. The solution *r* to the right of *B* is therefore constant and equal to the axon radius *R*_*a*_ = 1 (after nondimensionalisation). This implies

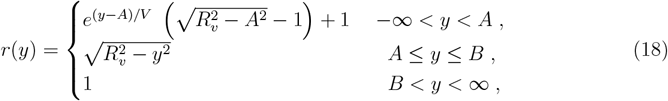

which still depends on *A*. The point *B* is to the right of *x* = 0 and defined by 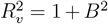, i.e.

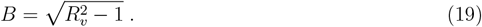

We finally use that *h*^*′*^(*A*) = *r*^*′*^(*A*) to compute *A* as a function of *V*. Using (18) this equation reads 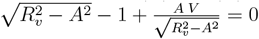 which implies that

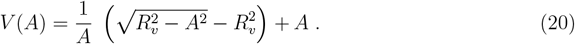

Note that the function 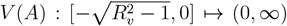 is monotonically increasing since 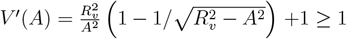 and convex since 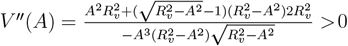 where we use 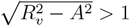.

In addition to 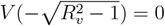 and lim_*A*→0_ *V* (*A*) = +∞ we also find that 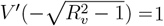 and lim_*A*→0_ *V*^*′*^(*A*) = +∞. As a consequence, taking the reciprocal values of the derivatives, we find that 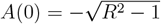, lim_*V*→∞_ *A*(*V*) = 0, *A*^*′*^(0) = 1 and lim_*V*→∞_ *A*^*′*^(*V*) = 0.

We also consider *A* as a function of *V*, namely the inverse of (20). As the inverse of *V* (*A*), it is monotonically increasing and concave. Note that in principle (20) can be inverted using the solution formulas for quartic equations. The result, however, is not of practical value, we therefore plot the graph of *A*(*V*) = *V*^−1^(*A*) numerically in figure 6.

**Figure 6:**
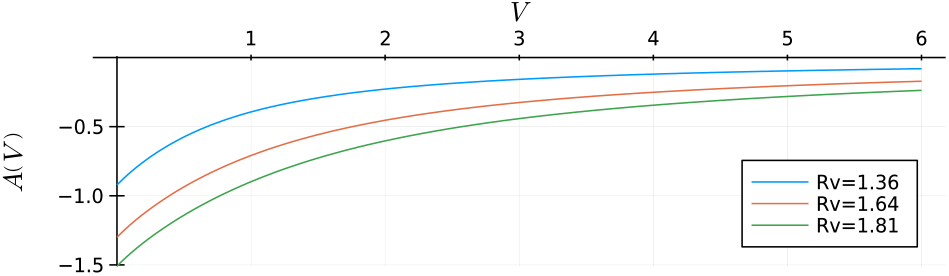
The position of the point *A* where the axon detaches from the vesicle as a function of the vesicle velocity *V* for various vesicle radii.

Finally, to find the vesicle speed *V*, we compute the density of repulsive force *λ* using the second equation of the system (17), which implies that *λ* = *r* – 1 − *V r*^*′*^. This implies that the Lagrange multiplier *λ* is given by

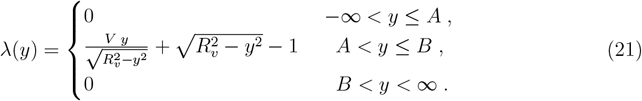

In figure 7 we show the profile of the interaction force *λ* as well as the traction *f* := *λ*(*y*)*h*^*′*^(−*y*). The traction is an odd function if *V* = 0 (figure 7A) and doesn’t generate any net traction on the vesicle since its integral in (16) vanishes. For *V >* 0 significant traction is generated along the front part of the vesicle (figure 7B).

**Figure 7:**
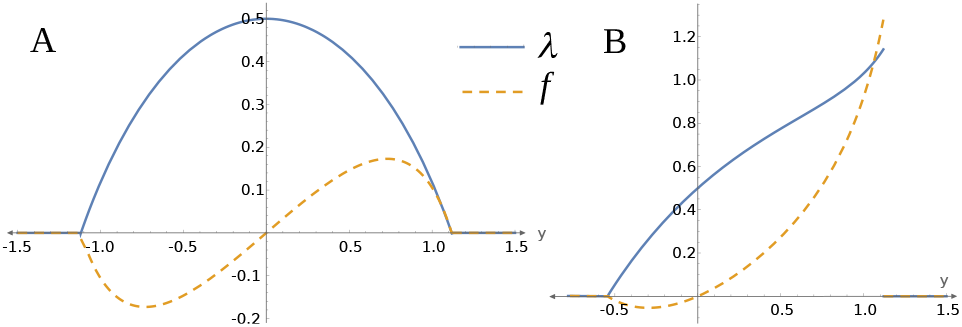
Interaction force *λ* and local traction *f* (*y*) = *λ*(*y*)*h*^*′*^(−*y*) for V=0 (A) and for the numerical example in figure 5 (B) where *V* = 1.02, *A* = −0.54, *B* = 1.12.

Note that as a consequence of (20) which is just *λ*(*A*) = 0 it holds that *λ >* 0 on (*A, B*] which confirms that same conclusion drawn from the constructive definition of *λ* in (11).

We substitute (21) in the first equation of (17) keeping in mind that *A* depends on *V* through the inverse of (20),

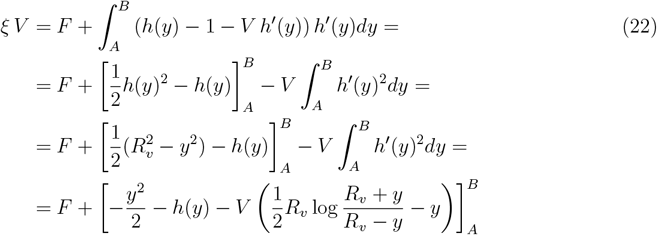

where we used that 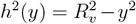 and the derivative of *h*^*′*^ which we obtain differentiating (7).

For the right endpoint of the interval [*A, B*] we substitute (19) and use (20) obtaining

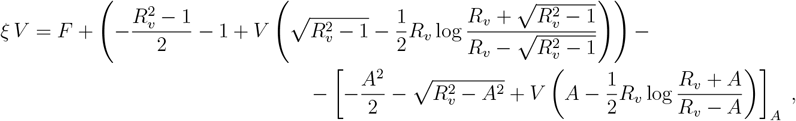

which, after rewriting, becomes

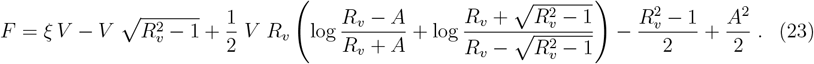

Note that *A* is taken here as a function of *V*, namely the inverse of (20). Since the solution formulas for quartic equations are unpractical, we plot the graph of (23) numerically (see figure 8) interpreting it as force-velocity relation for the axonal transport of vesicles with (scaled) radius *R*_*v*_.

**Figure 8:**
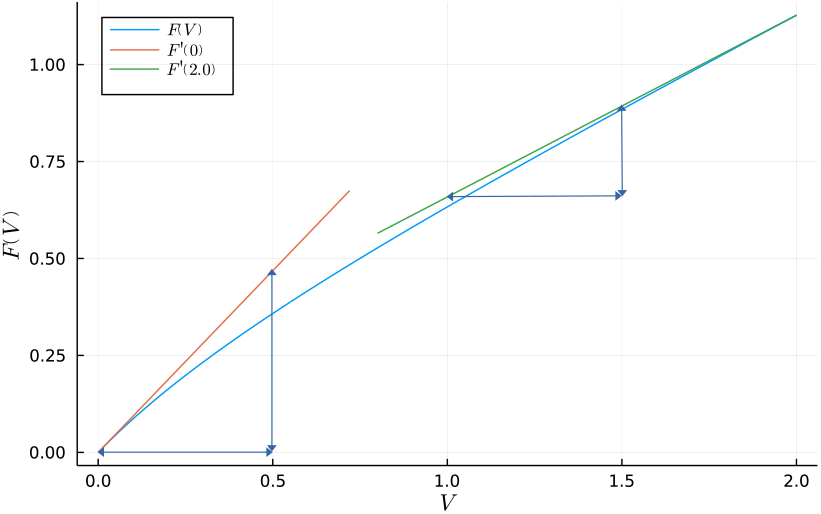
Force-velocity relation for *R*_*v*_ = 1.64 and *ξ* = 0.

In the supplementary section B we confirm the validity of the equation (23) numerically.

Note that *A*(0) = −*B* and therefore *F* (0) = 0. Taking derivatives in (22) and (23) we obtain the following expressions for the derivative *F*^*′*^(*V*) which is the marginal force to accelerate the vesicle (keep in mind that this is a model for over-damped dynamics),

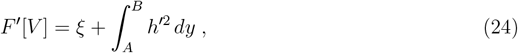

which implies that *F* is monotonically increasing since *F*^*′*^ *>* 0 and it is also concave since *F*^*′′*^(*V*) = −*h*^*′*^(*A*)^2^ *<* 0. Substituting 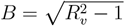 we obtain

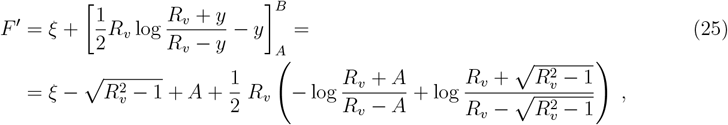

which can only be evaluated numerically. At the endpoints, i.e. for *V* = 0 and in the limit where *V* → ∞ we can find, though, explicit values using 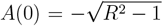 and lim_*V*→∞_ *A*(*V*) = 0, namely

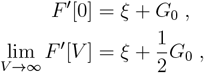

where

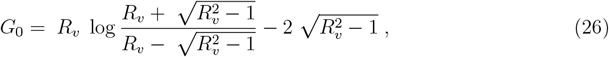

which is non-negative because *F*^*′*^ *>* 0 for all *ξ* ≥ 0. Not taking into account linear drag with coefficient *ξ* exerted on the vesicle through the crowded cytoplasm, the marginal friction *F*^*′*^[*V*] decreases by a factor 2 as the velocity increases (figure 8).

## 5 Vesicle size dependence of transport speed

In this section, it is our goal to understand how the vesicle speed predicted by our model depends on the vesicle size *R*_*v*_. We will evaluate this dependence numerically and use asymptotic expansion to compute an approximate solution when the vesicle radius *R*_*v*_ is similar to the axon radius *R*_*a*_. Note that after non-dimensionalisation the axon radius is 1. To facilitate the asymptotic analysis we therefore write the vesicle radius as *R*_*v*_ = 1 + *ε*^2^ and consider *V*_*ε*_ as the solution of the system (23), (20). Numerical solutions of this system for fixed parameters *F* and *ξ* illustrate that the predicted vesicle speed decreases with larger vesicle diameter (figure 9) in line with experimental observations[11].

**Figure 9:**
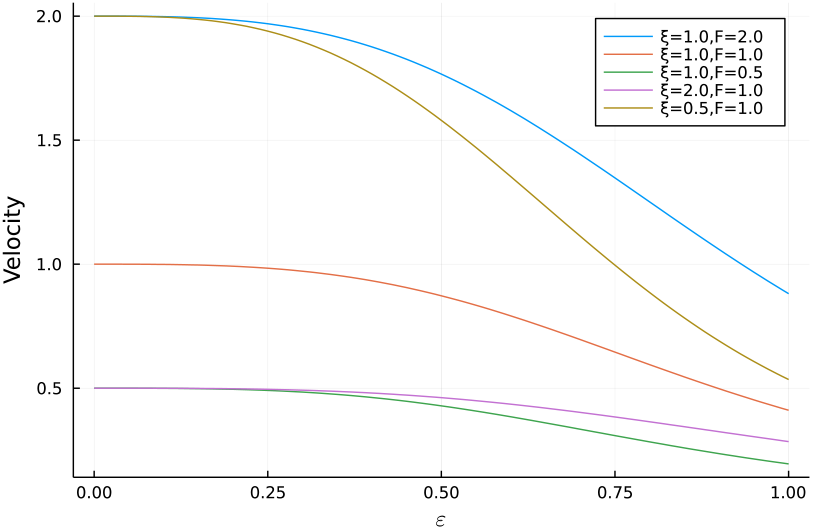
The velocity decreases as the cargo diameter, represented by 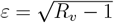, increases. Compared to the reference graph in red, the blue and green graphs correspond to fixed *ξ* with different values of force *F*, while the pink and red graphs represent fixed force *F* with various values for (scaled) drag friction *ξ*.

For the asymptotic analysis we start with the interface points where the axon wall detaches from the vesicle. In the modified notation the interface point in front of the vesicle is at

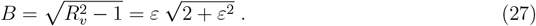

The interface point at the rear of the vesicle, on the other hand, is determined by equation (20) which relates it to the velocity *V*. Using the modified notation, this equation reads

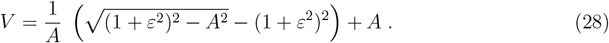

Using the ansatz *A* ∼ *A*_1_*ε* + *A*_2_*ε*^2^ + … we find that the expansion up to order 5 in *ε* is given by

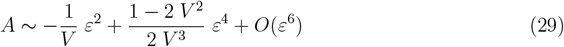

Note that this expansion is not valid in regimes where *V* is small as it yields un-physically small or large values of the position *A* where the vesicle detaches from the axon.

We now turn to equation (23) to compute an approximate solution for the vesicle transport velocity. We couple this equation to the expansion (29) and use the ansatz *V* ∼ *V*_0_ + *V*_1_*ε* + *V*_2_*ε*^2^ + *V*_3_*ε*^3^ + …. Equating coefficients of equal powers of *ε* yields the asymptotic solution (detailed in supplementary section A)

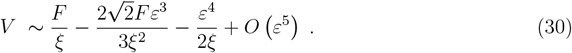

In the same way as the expansion of *A* in (29), the expansion of *V* is not valid when *ξ* is small in which case the expansion would predict negative velocity. In figure 10 we compare the numerical solution for the vesicle velocity to the asymptotic approximation given by the leading order terms up to order 3 in (30).

**Figure 10:**
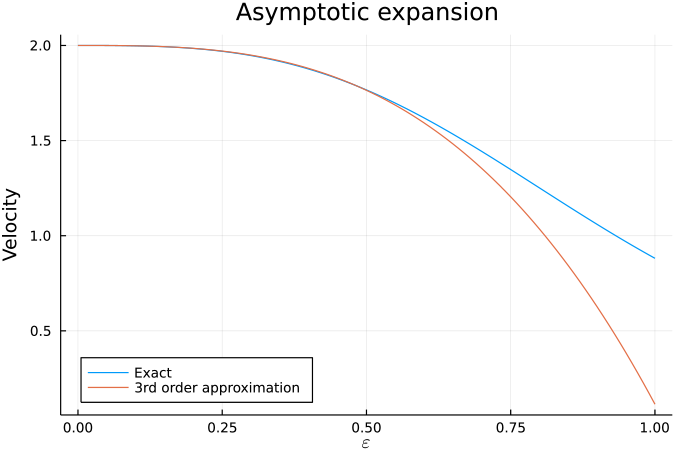
Comparison of the numerically computed graph of *V* as a function of 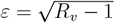 representing vesicle radius with the asymptotic expansion of *V* in (30) up to 3rd order in *ε* for *ξ* = 1, *F* = 2.

Finally we rewrite the asymptotic solution (30) up to third order in *ε*, 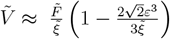, in dimensional form substituting the definitions of the dimensionless independent and dependent variables, 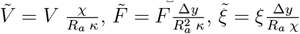 and 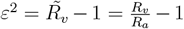 from section 3. We find that

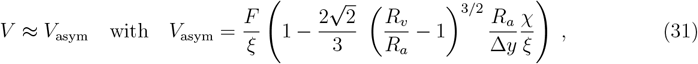

which quantifies the decrease of transport speed as a function of vesicle radius *R*_*v*_ in the regime when the vesicle radius is close to the axon radius and drag friction is not too small.

Graphs of *V*_asym_ are shown in figure 11. The left figure shows the decrease of *V*_asym_ as a function of the cargo radius *R*_*v*_ for fixed axon radius *R*_*a*_ (Table 1). Note that the predicted 40% decrease of the velocity as the cargo exceeds the axon radius by 50% is consistent with the decrease in vesicle speed observed in retrograde axon trafficking in both proximal and distal axon segments of DIV14 rat hippocampal neurons [11](figures 7b and 7c). We also remark that the magnitude of the decrease in the asymptotic velocity (31) visualised as black arrows in figure 11 scales linearly with the ratio of actomyosin ring viscosity *χ* and positional drag *ξ*.

**Figure 11:**
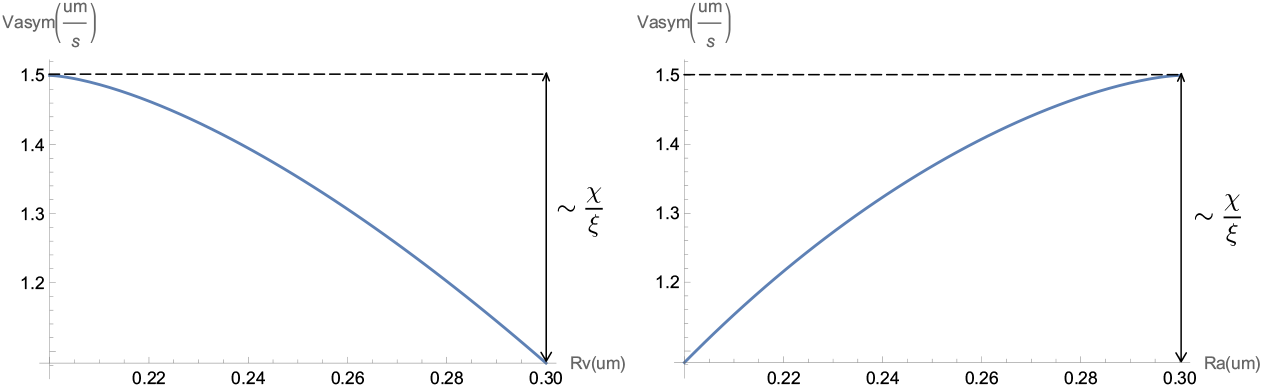
Left: Decrease of *V*_asym_ as a function of cargo radius *R*_*v*_ for fixed axon radius *R*_*a*_ (with parameters according to Table 1). Right: Increase of *V*_asym_ as a function of axon radius *R*_*a*_ for fixed cargo radius *R*_*v*_ (parameters according to Table 1).

The right graph in figure 11 shows the increase of *V*_asym_ as a function of the axon radius *R*_*a*_ for fixed cargo radius *R*_*v*_ (with parameters according to Table 1). We remark that inhibition of myosin-II motor proteins is correlated with increased axon diameter 2*R*_*a*_ as reported in [11](figure 4d: ctrl: 0.35 μm, myosin-II inhibition: 0.55 μm which amounts to an increase of roughly 50%). Our model therefore predicts an increase of cargo velocity upon myosin inhibition of about 40% which is in line with experimentally observed cargo velocities in the proximal segment of neuron axons (figure 7b in [11]).

## 6 Discussion

In this study, we establish a mathematical model for the interaction of the membrane associated periodic skeleton (MPS) in axons with large cargo vesicles undergoing axonal transport. The model accounts for transient swelling caused by the excess size of the cargo and for the mechanical resistance this imposes on its motion. The resulting model is a free boundary problem and resembles the classical obstacle problem.

There are several limitations to our model since it is constructed as a minimal model for the interaction of the cargo vesicle with the axon wall. One significant simplification is that we treat the vesicle as a solid sphere while in reality, it is viscoelastic with different mechanical properties of vesicle membrane on the one hand and vesicle lumen on the other hand (see sketch in figure 1). Deformation of large vesicles would likely reduce the resistance to vesicle motion through axon wall resistance and through friction with the axon wall and the cytoplasm. While the effect would be most pronounced for very large vesicles, in this study we focus on the regime when *ε* is small.

Another question which we neglect in this study concerns the flow of cytoplasm caused by the translocation of large cargo vesicles. We believe that compensating fluxes should be present in the narrow margins between large cargo vesicles and the axon membrane as sketched in figure 1 and visible in the electron micrographs in figure 2 of [11]. In our model, the resistance of the cytoplasm to large cargo movement is captured by the drag friction with coefficient *ξ*. A more detailed treatment of the cytoplasm would require coupling to a fluid model which we postpone to future studies.

We consider quasi-stationary solutions and reformulate them as force-velocity relation for the transport speed of the cargo. It predicts that when neglecting drag, the marginal force - the slope of the force-velocity relation - decays by a factor of 2 starting from when the vesicle is at rest, which can be tested experimentally.

We also compute - numerically and asymptotically as closed-form expression - the size dependence of the transport speed. It predicts that the velocity decays for larger vesicles which is in line with recent observations [11]. It also predicts that the transport speed as a function of vesicle size follows a power law with exponent 3*/*2 which is amenable to experimental verification.

Other predictions can be obtained from the dimensional expression of the asymptotic cargo velocity (31). The ratio of actomyosin ring viscosity *χ* and positional drag *ξ* governs the magnitude of the decrease in the asymptotic velocity visualised as black arrows in figure 11.

It is striking that the stiffness of actomyosin rings *κ* is not present in the asymptotic expression (31). We believe that this is because at leading order, i.e. for not too large vesicles, the resistance of stiff transversal actomyosin rings balances the support they provide at the back. The dimensional reformulation of the fourth-order term in (30), however, is given by −*κ* (*R*_*v*_ − *R*_*a*_)^2^*/*(2 *ξ* Δ*y*) and includes *κ* indicating that particularly for rather large vesicles transversal ring stiffness does reduce cargo velocity.

The particular way in which these quantities, transversal ring viscosity *χ* and stiffness *κ*, affect the cargo’s velocity could potentially be exploited to develop techniques to control the transport of particularly large cargo vesicles in axons.

In its present form, the model is designed to be a foundation for future computational studies of axonal transport through stochastic simulation of molecular motors dynein and kinesin involved in cargo transport. Some additional aspects of the mechanics involved in the axonal transport of large vesicles will most likely be of interest and should therefore be incorporated into the model. One is friction between the axon wall and vesicles taking into account both normal forces and shear forces which has already been suggested as a potential explanation of the size-dependence of the vesicle speed [11]. Such friction could be either caused by direct interaction or mediated by cytoplasm viscosity [18]. Another striking aspect of axonal mechanics is the coupling between neighbouring actin rings which is mediated by spectrin and sustains axial tension [25], therefore coupling radial and axial tension [26]. Incorporating spectrin into the present model would most likely introduce 2nd order derivatives into the continuum model (16), lead to different free boundary conditions and require a fundamentally different mathematical treatment due to the presence of the second order derivative.

## Acknowledgements

The authors acknowledge funding from the Australian Research Council (ARC) Discovery Program (grant number DP180102956), awarded to D. B. O. An RTP scholarship funded by the University of Queensland (UQ), awarded to N. R.

## A Asymptotic expansion of transport velocity

Starting with the first equation in (22), we have the following integral equation

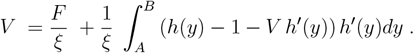

Now we substitute *h* and its derivative given in (7) obtaining

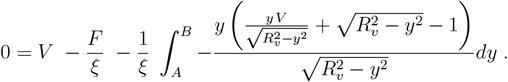

Coupling this to the asymptotic expansions of *A* and *B* given in (29) and (27) we conclude that

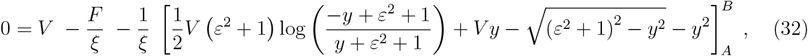

where we used *R*_*v*_ = 1 + *ε*^2^. Finally, substitute the asymptotic expansion for *V* into (32),

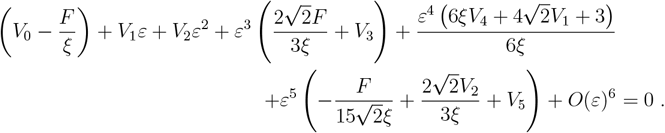

Equating coefficients of equal powers of *ε* we obtain

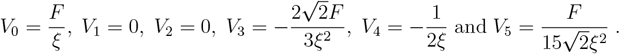

## B Numerical test of quasi-steady state approximation

In this supplementary section we numerically test the validity of the quasi-steady state approximation (23). To this end we solve the time discrete model (8)-(9) numerically, extract the cargo velocity and compare it to the result obtained from numerically solving (23). The numerical results coincide (figure 12).

**Figure 12:**
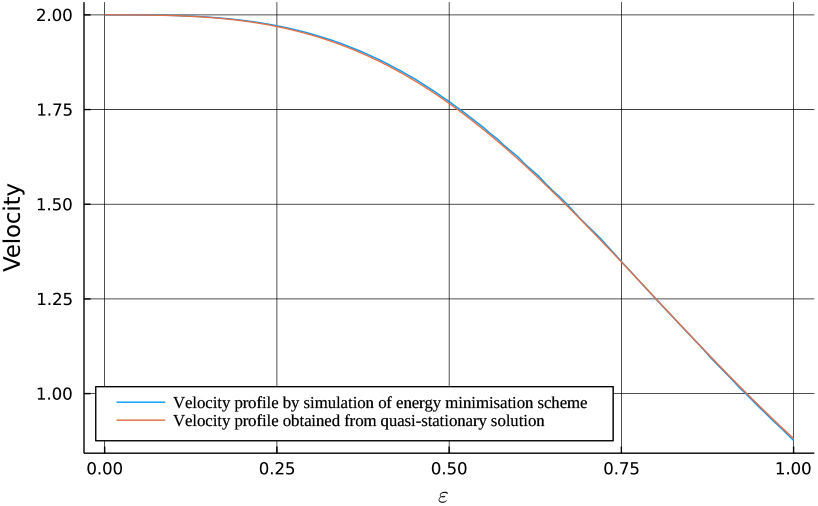
The velocity obtained through solving the discrete system (8)-(9) for different values of 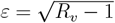 coincides with the velocity computed numerically using the quasi-steady state approximation of the continuum model (23) for *ξ* = 1, *F* = 2.

